# Epithelial state-transitions permit inflammation-induced tumorigenesis

**DOI:** 10.1101/2025.06.22.660911

**Authors:** Edward J Jarman, Anabel Martinez Lyons, Yuelin Yao, Aleksandra Rozyczko, Scott H Waddell, Andreea Gradinaru, Paula Olaizola, Kyle Davies, Rachel V Guest, Stephanie Röessler, Timothy J Kendall, Owen J Sansom, Ava Khamseh, Luke Boulter

## Abstract

Chronic inflammation across tissues is associated with an increased risk of developing cancer^1–3^. While potentially oncogenic somatic mutations have been demonstrated to persist and expand in healthy organs^4–6^, what triggers a subset of cells harbouring deleterious mutations to transition into a neoplasm or an aggressive adenoma with poor prognosis^7,8^ is not well-understood. Unlike normal, healthy cells, benign cells harbouring mutations perceive inflammation in chronic disease differently, potentiating the progression from physiological inflammation to tumorigenesis^9^. Here, we reveal that a subset of epithelial cells with mutations are poised to transition from pre-neoplastic state to early neoplasm, through rewiring of epithelial IL-1β responses and inflammatory macrophage recruitment. We characterise this process by leveraging a mouse model of biliary tract cancer (cholangiocarcinoma), in which deleterious mutations are introduced to tumour suppressor genes in common cancer pathways (*Trp53* and *Pten*), and by quantifying differences in cell states and corresponding gene expression dependencies in the absence or presence of liver inflammation. Critically, we find that targeting the epithelial-derived signals of tissue-wide inflammation (namely COX2) is insufficient to limit tumorigenesis; rather, targeting the reactivation of oncogene-induced developmental signals, such as NOTCH, prevents this pre-neoplastic to neoplastic transition, demonstrating that oncofoetal switching is a pharmacologically-tractable target in patients with a high risk of developing cancers on the background of inflammation.

## Main

As organisms age, they accumulate mutant cells across their lifetime; indeed, such mutant cells exist at surprisingly high frequencies in otherwise “healthy” somatic tissues^5,6,10,11^. Despite containing large numbers of mutant cells in a range of organs, the actual instance of malignant transformation within a single organism is relatively rare, suggesting that mutation alone is often insufficient to reprogramme cell plasticity to initiate cancer^12^, but can promote clonal selection^6,13^. A major risk factor for the development of cancer across organs is chronic inflammation^2,9^, yet how this essential pathophysiological wound-healing response intersects with the accumulation of otherwise benign mutant cells to drive cell state plasticity, which permits carcinogenesis, is unclear.

In the liver, the risk of developing biliary tract cancers, including cholangiocarcinoma (CCA), is substantially increased in patients with chronic inflammatory conditions such as liver fluke infection, primary sclerosing cholangitis (PSC) and intrahepatic gallstones^14^ – despite these well-defined risk factors, the sequence of cellular and molecular events that lead normal biliary epithelial cells (BECs) to transition into dysplasia and subsequently neoplasia is poorly understood. BECs demonstrate both anatomical and functional heterogeneity^15^ ranging from smaller, mono-ciliated columnar cells which line the terminal ductules of the biliary tree to larger cells with immunoregulatory and secretory roles within the larger ducts (Fig. 1a). Different inflammatory aetiologies of CCA show tendencies to affect different anatomical locations across the biliary tree^16^ and in population sequencing studies CCA of different anatomical origins demonstrate diverse mutational patterns^17–19^, suggesting a hitherto unappreciated interplay between epithelial cell state, inflammation and the mutational patterns which synergise for neoplastic growth.

**Figure 1:**
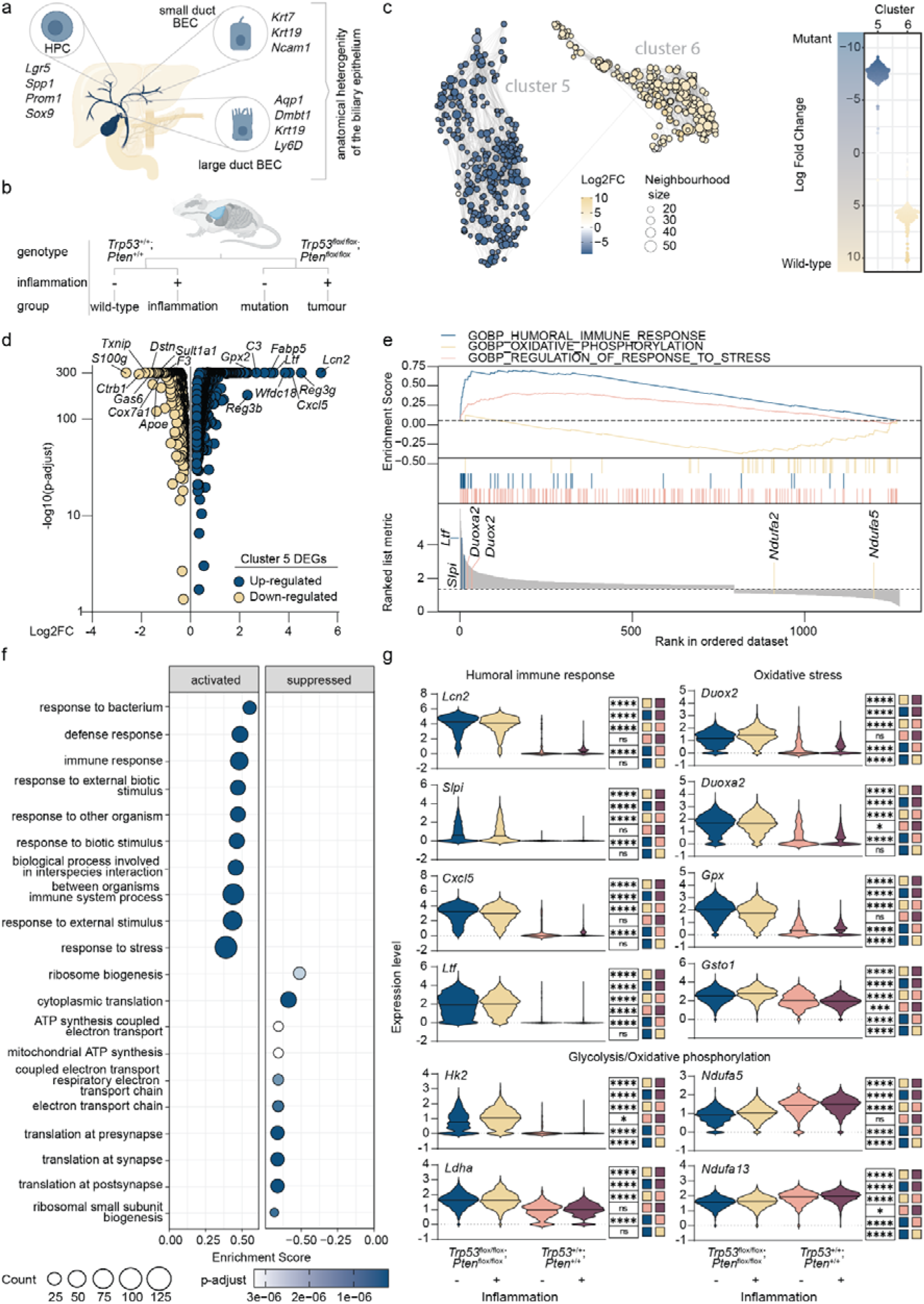
Large duct BECs are uniquely susceptible to transformation: **a.** Schematic demonstrating the anatomical and cellular heterogeneity found within the mammalian biliary tree and **b.** demonstrating the experimental combinations in this study. **c.** Differential abundance testing in large duct BECs from *Trp53*;*Pten*-mutant and wild-type animals showing neighbourhoods identified in clusters 5 (blue) and 6 (beige), left panel. Swarm plot showing relative abundance differences between neighbourhoods in clusters 5 and 6, right panel. **d.** Volcano plot demonstrating the up (blue) and down (beige) regulated genes in cluster 5 compared to cluster 6. **e.** Gene Set Enrichment Analysis of *Trp53*;*Pten*-mutant BECs. **f.** Top 10 activated and suppressed GOTerms in *Trp53*;*Pten*-mutant large duct BECs compared to wild-type counterparts. **g.** Histograms showing mRNA expression of genes involved in humoral immune signature, oxidative stress and glycolysis/oxidative phosphorylation in either *Trp53*;*Pten*-mutant or wild-type BECs with or without inflammation.

Here, we show that heterogenous epithelial cells within the bile duct differentially respond to common cancer pathway mutations by modifying their response to tissue-wide inflammation, promoting specific subsets of BECs to enter a primed, pre-neoplastic cell state. By mapping the molecular route through which pre-neoplastic BECs transition into a neoplasm, we identified that persistent inflammation drives an IL-1β-NOTCH signalling axis, supporting the malignant transformation of large-duct BECs, which critically, when restricted, limits neoplastic growth. While the accumulation of mutant cells within tissues is inevitable, our data demonstrates that physiological inflammation is maladaptively interpreted by these mutant cells and strongly supports the development of prophylactic treatment strategies for patients with chronic inflammatory disease and an elevated risk of biliary tract cancer.

## Large duct epithelial cells are uniquely susceptible to pre-neoplastic transformation

Mutations in the PTEN/PI3K and P53 pathways are relatively common in both established CCA^7^ and pre-malignant tissue from patients with primary sclerosing cholangitis (PSC) or biliary intraepithelial neoplasia (BilIN)^20^. Given the intrinsic cellular heterogeneity of cells within the bile duct (Fig. 1a), we sought to dissect how heterogenous biliary populations respond to common cancer-associated genetic mutations by leveraging a murine model of CCA^21^ in which *Pten* and *Trp53* are specifically inactivated in BECs (while simultaneously and irreversibly labelling transformed BECs with tdTomato, following recombination with *Krt19*-CreERT, herein called *Trp53*;*Pten*-mutant, Fig. 1b and Extended Data Fig. 1a).

Nascent tdTomato-positive, *Trp53*;*Pten*-mutant cells fail to spontaneously form tumours, however following treatment with the liver inflammatory agent thioacetamide (TAA)^21,22^, tdTomato lineage traced cells form histological neoplasms within three weeks (Extended Data Fig. 1b). Single cell RNA (scRNA) sequencing of 21,372 lineage-traced BECs isolated from *Trp53*;*Pten*-mutant and control mice (in which *Trp53* and *Pten* are intact) with and without inflammation revealed seven principal populations of BECs. Based on canonical marker expression (Supplementary Table 1), we characterised these as small duct BECs (clusters 1-3), large duct BECs (clusters 5 and 6), BECs undergoing ductular remodelling (cluster 4) and a proliferative cluster (cluster 7), the latter two of which only arise when inflammation is present (Extended Data Fig. 1c and 1d). Using Milo differential abundance testing^23^ (Methods) we identified distinct cellular neighbourhoods enriched in mice exposed to inflammation alone versus those with only *Trp53*;*Pten*-loss without exposure to inflammation (Extended Data Fig. 2a), thereby confirming that inflammation *per se* supports the emergence of both proliferative and regenerative cell clusters that are enriched in genes associated with oxidative stress and respiratory processes (Extended Data Fig. 2b and 2c); and these populations do not arise in the absence of inflammation. Furthermore, equivalent neighbourhood testing in cells with tumour suppressor loss highlights a clear positive enrichment in cluster 5, comprising large duct BECs. Moreover, GO analysis of terms associated with *Trp53*;*Pten*-deletion highlight this loss as an unexpected driver of epithelial immune regulation, including terms associated with direct response to bacterial pathogens and dysregulation of the immunoproteasome, which drives antigen production for the antigen presentation machinery (Extended Data Fig. 2b and 2d, Supplementary Table 2).

While top-level transcriptional analysis of isolated BECs showed that cluster 5 is enriched for *Trp53*;*Pten* mutant cells, cluster 6 is enriched for wild-type large duct BECs. This mutual exclusivity is independent of inflammation (Fig. 1c and 1d, Supplementary Table 3), suggesting that large duct BECs are more sensitive to mutational perturbation when compared to small duct BECs, which do not transcriptionally segregate based on mutational status. Gene set enrichment analysis between clusters 5 and 6 shows that *Trp53*;*Pten*-loss promotes a stress response signature driving anti-microbial and humoral immune signatures, as well as genes associated with cancer-associated metabolic reprograming (Fig. 1e-g), leading us to hypothesise that mutant large duct BECs alter their transcriptional phenotype to enter a transformed cellular state, which is poised for inflammation-induced neoplastic transition.

To further define whether this transformed state is a *bona fide* neoplastic pre-cursor, we focused on identifying cellular neighbourhoods which only arise when *Trp53*;*Pten* are lost and inflammation is present and identified a unique neighbourhood within cluster 4 (Fig. 2a). GO term analysis of DEGs within this specific neighbourhood were enriched for epithelial immune processes akin to mutant large duct BECs in cluster 5 (Fig. 2b, Extended Data Fig. 3a and Supplementary Table 4) and using a combined gene expression matrix of the top 50 upregulated genes we found a distinct overlap in gene expression between *Trp53*;*Pten*-mutant large duct BECs and neoplastic cells (Fig. 2c) suggesting that they are the immediate progeny of transformed large duct BECs.

**Figure 2:**
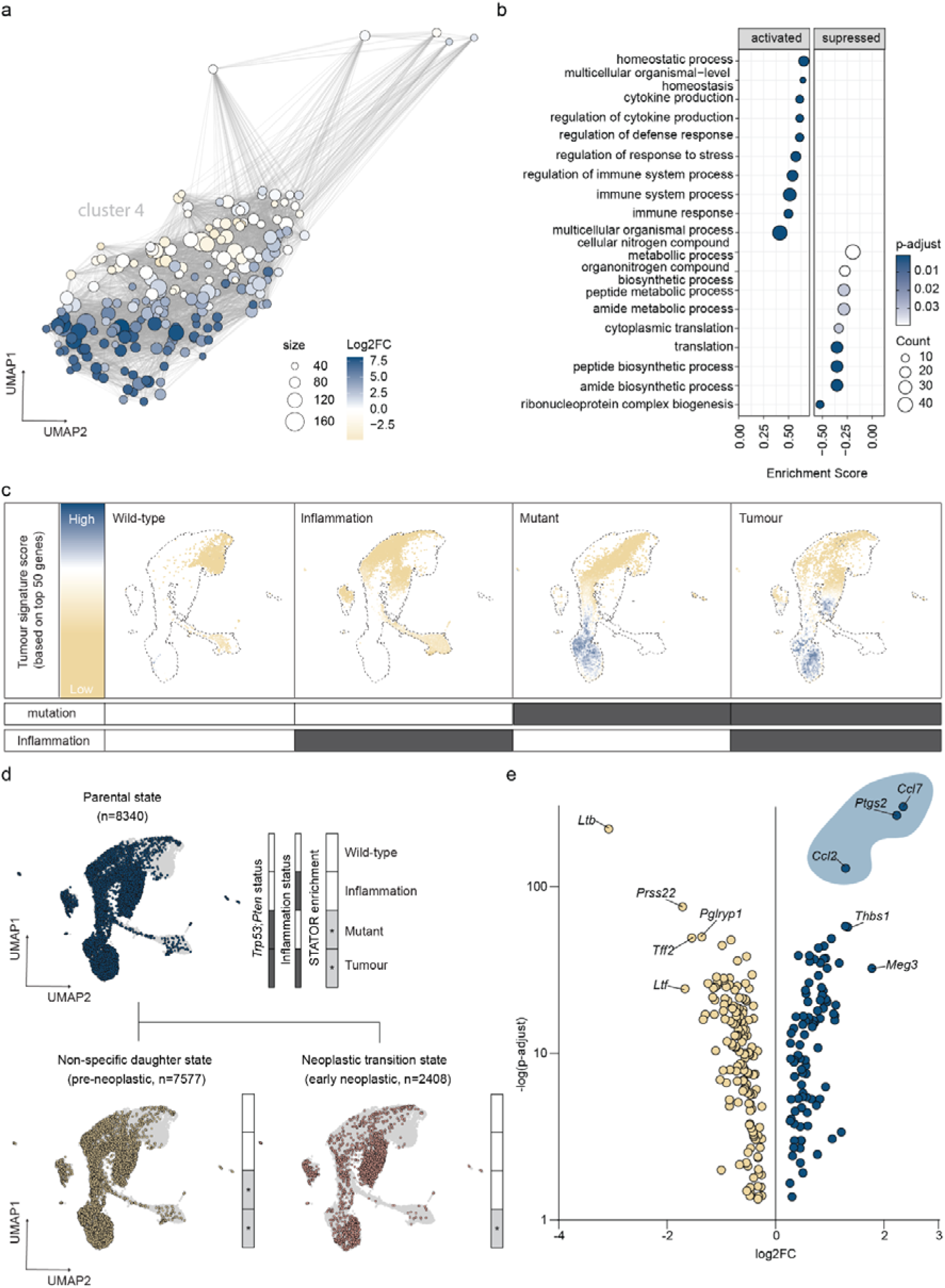
Transcriptional co-dependencies identify the earliest moments of neoplastic formation: **a.** Differential abundance testing (with Milo) of cluster 4, highlighting a group of neighbourhoods which only arise in *Trp53*;*Pten*-mutant animals when inflammation is present (blue). **b.** Top 10 activated and suppressed GOTerms in neighbourhoods identified in *Trp53*;*Pten*-mutant, inflamed neighbourhoods. **c.** UMAP projection of the top 50 differentially expressed genes identified through differential abundance testing (Methods) in *Trp53*;*Pten*-mutant, inflamed neighbourhoods in untreated, wild-type animals (wild-type), genetically wild-type animals with inflammation (inflammation), transgenic animals with BEC *Trp53*;*Pten*-loss (mutant) and animals with BEC *Trp53*;*Pten*-loss and concurrent inflammation (tumour). Blue shows high expression and beige low expression. **d.** Stator cell states imposed on UMAP showing parental state (blue) and cells with BEC *Trp53*;*Pten*-loss, exposed to inflammation that do not (beige, pre-neoplastic) and do (pink, early-neoplastic) become neoplastic based on probabilistic gene expression dependencies. Legend: dark grey bars denote experimental conditions; light grey define the state(s) present within those cells. **e.** Volcano plot showing differential gene expression between pre-neoplastic and early-neoplastic cells with a log2 fold change >0.25 and an adjusted p<0.05 corrected using a Benjamini-Hochberg for multiple testing.

## Probabilistic gene expression dependencies define the emergence of neoplastic cell states

Not all mutant cells become neoplasms even in the presence of inflammation. We sought to separate the cell states of *Trp53*;*Pten* mutant cells from those becoming neoplastic using Stator (Methods), which identifies data-driven novel cell states based on higher-order gene expression dependencies^24^. Stator is entirely agnostic to experimental grouping and identified unique cellular states that only emerge when *Trp53*;*Pten*-mutant cells are exposed to inflammation (Extended Data Fig. 3b). Stator’s hierarchical clustering of higher-order gene dependencies identified the branching point at which cellular states become specific to neoplasms thereby allowing for the direct comparison between *Trp53*;*Pten*-mutant cells which do (early neoplastic) and do not (pre-neoplastic) contribute to neoplasia (Fig. 2d). These early-neoplastic states are defined by both *Ptgs2*/*Ccl2*/*Mmp7* and *Ccl7*/*Mmp7*/*Ccl2* gene expression modules (Fig. 2e and Supplementary Table 5).

## COX2-positive cellular states define human biliary neoplasms and are present in high-risk patient groups

Recent work identified COX2/PGE_2_ production by cancer cells as part of a paracrine feedback loop by which macrophages drive IL-1β signalling in pancreatic ductular adenocarcinoma (PDAC), a cancer which shares many histopathological features with CCA^9,25^. Given we have shown that early neoplasms are defined by the co-expression of *Ptgs2* (COX2) and monocyte recruitment factors *Ccl2* and *Ccl7*, we wanted to determine whether tumour initiation in the bile duct occurs through an analogous process to that described in PDAC. Indeed, COX2-positive, *Trp53*;*Pten*-mutant early neoplasms form intimate associations with IL-1β-positive, F4/80-positive macrophages (Fig. 3a) and pre- and early-neoplastic Stator states demonstrate increased transcriptional response to IL-1β (Fig. 3b, Supplementary Table 6).

**Figure 3:**
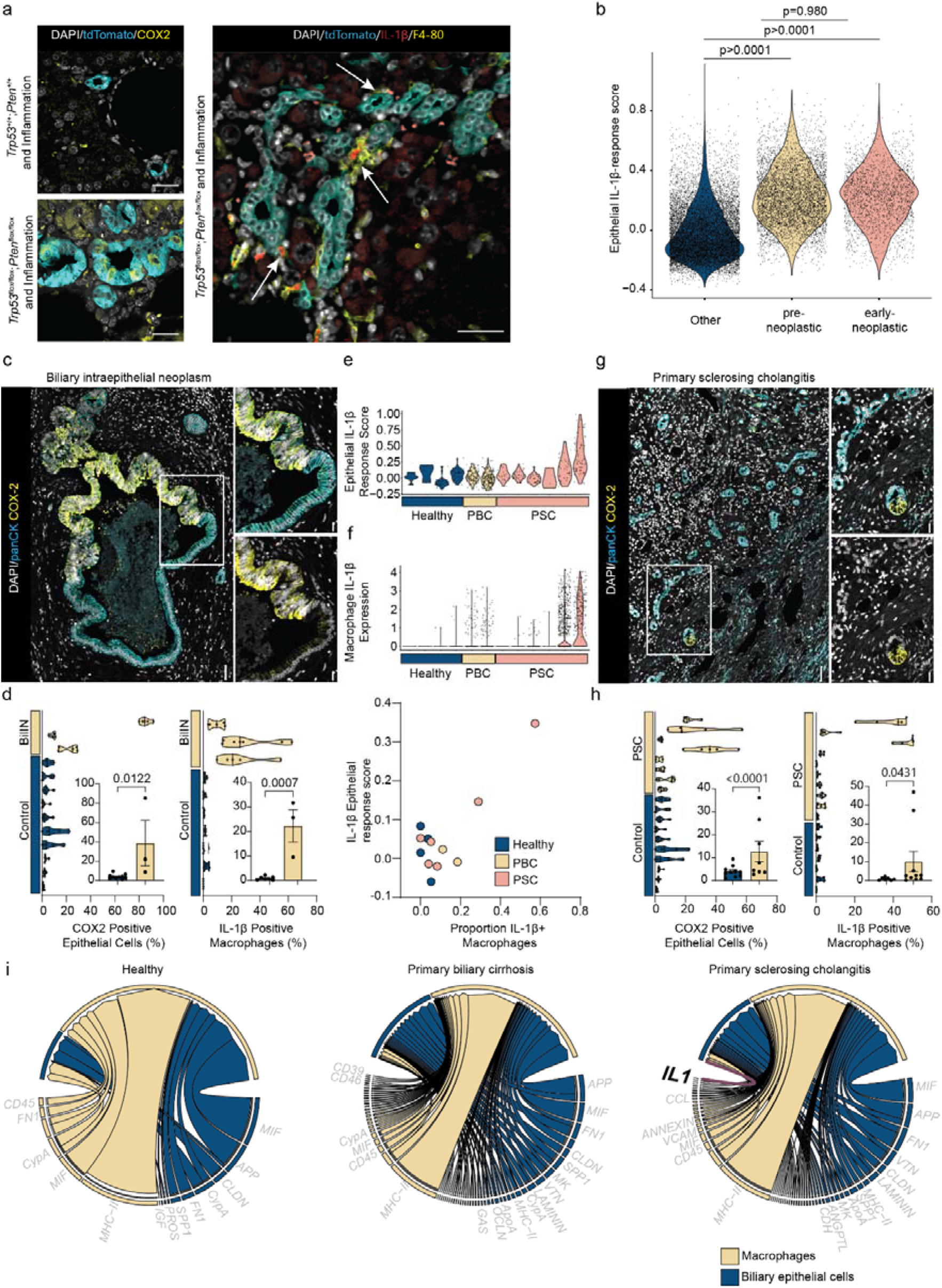
Pre-neoplastic cell states are common from mouse to human. **a.** Immunofluorescent staining of tdTomato positive BECs (cyan), COX2 (yellow) and DNA (white) in *Krt19*-CreER^T^;R26^RLSL-tdTomato^ animals with *Trp53*^flox/flox^;*Pten*^flox/flox^ (mutant) or *Trp53*^+/+^;*Pten*^+/+^ (wild-type) BEC, left panels. Right, *Krt19*-CreER^T^;R26^RLSL-tdTomato^;*Trp53*^flox/flox^;*Pten*^flox/flox^ neoplasms immunostained for tdTomato (cyan), IL1β (magenta) and F4/80 macrophages (yellow). Arrows denote IL1β-positive macrophages adjacent to tdTomato positive cells. Scale bar = 100µm. **b.** Aggregated expression matrix of an IL-1β transcriptional signature showing enrichment in pre-neoplastic (beige) and early-neoplastic (pink) compared to all other cells within the dataset (other, blue). **c.** Immunofluorescent staining of human BilIN for BECs (pan-cytokeratin, cyan) and COX2 (yellow), DNA (grey). Scale bar = 50µm. **d.** Quantification of COX2-positive epithelial cells, left graph and IL-1β-positive macrophages, right graph in BilIN compared to normal tissue. **e.** Expression of an epithelial IL-1β signalling response score in scRNA data from Andrews et al. from normal patients (blue), PBC (beige) and PSC (pink) patients**. f**. Expression of IL-1β in macrophages from normal patients (blue), PBC (beige) and PSC (pink) patients (upper panel). Correlation of epithelial IL-1β response and IL-1β macrophage expression in normal patients (blue), PBC (beige) and PSC (pink) patients (lower panel). **g**. Immunofluorescent staining of PSC tissue for BECs (pan-cytokeratin, cyan) and COX2 (yellow), DNA (grey). Scale bar = 50µm. **h.** Quantification of COX2-positive epithelial cells, left graph and IL-1β-positive macrophages, right graph in PSC compared to normal tissue. **i.** Ligand-Receptor interaction analysis between BECs and Macrophages in normal, PBC and PSC scRNA data derived from *Andrews, et al*.

In humans, biliary intraepithelial neoplasms (BilIN) arise within the intrahepatic ducts and gallbladder^26,27^ and while histologically analogous to pancreatic intraepithelial neoplasms, little is known about the molecular changes that promote BilIN formation. To determine whether COX2-expressing epithelial cells identified in mouse during neoplastic transition also exist in human disease, we immunostained for COX2-positive epithelial cells (demarcated using pan-cytokeratin) and IL-1β-positive macrophages in human BilIN tissue (Fig.3c). COX2 expressing epithelial cells formed tight boundaries between areas of dysplasia and normal ductular tissue; moreover, they were only detected in BilIN but not in healthy ducts (Fig. 3c). Similar to mouse neoplasms, IL-1β expressing CD68-positive macrophages were also highly present in areas adjacent to neoplastic disease (Fig. 3d).

While BilIN are considered early neoplastic lesions for CCA, they are rare. Patients with primary sclerosing cholangitis (PSC) have a ∼7-13% lifetime risk of developing CCA and represent the most high-risk patient group in the West^14^. We reason then, that a subset of PSC patients would harbour COX2 positive pre-neoplastic cells. Using publicly available scRNA-seq data of human PSC, primary biliary cholangitis (PBC) and healthy liver^28^ we sought to define whether neoplastic cell states were present in these patients. Whilst PSC is considered an idiopathic inflammatory disease predominantly affecting larger bile ducts, PBC is limited to smaller, interlobular intrahepatic ducts and does not have a strong associated risk of developing CCA^8^. Indeed, patients with small duct PSC also have a low incidence of CCA development^29,30^. Using a gene signature to define responsiveness to IL-1β signalling^9^, we only identified BECs with this ability from patients with PSC, not PBC or healthy tissue (Fig. 3e, Extended Data Fig. 4 and Supplementary Table 6), which additionally correlates with the expression of IL-1β by macrophages (Fig. 3f). Critically, this IL-1β-epithelial signature is not found in BECs from healthy or PBC patient samples even when IL-1β-positive macrophages are present, implying that a subpopulation of BECs in PSC patients are specifically responsive to macrophage-derived IL-1β. Using an independent, expanded group of PSC patients we found COX2-positive BECs in a subgroup of samples, showing that the emergence of COX2-positive cells and the presence of IL-1β macrophages is not a universal feature of PSC, but is restricted to specific patients (Fig. 3g and 3h). Within PSC patients, ligand-receptor interaction modelling between epithelial and macrophage clusters from healthy, PBC and PSC scRNA-seq data only identifies IL-1β signalling from macrophages to the IL-1R/IL-1RAP complex in epithelial cells in PSC (Fig. 3i, Supplementary Table 7).

## Inflammatory context defines therapeutic efficacy of COX2 inhibition

The presence of COX2-expressing pre-neoplastic BECs in PSC patients provides a potential target to prevent the development of CCA in these high-risk patients. In PDAC, inhibition of COX2 limits tumour growth through reduced IL-1β-positive macrophage recruitment^9^. Therefore, we inhibited COX2 during inflammation-driven tumour-initiation using either low dose aspirin, which inhibits COX1 and COX2 or Celecoxib, a selective antagonist of COX2^31,32^. Neither aspirin or Celecoxib had an inhibitory effect on tumour initiation and growth in our model (Fig. 4a), suggesting that the wider inflammatory context in chronic ductular disease overcomes neoplastic inflammation.

**Figure 4:**
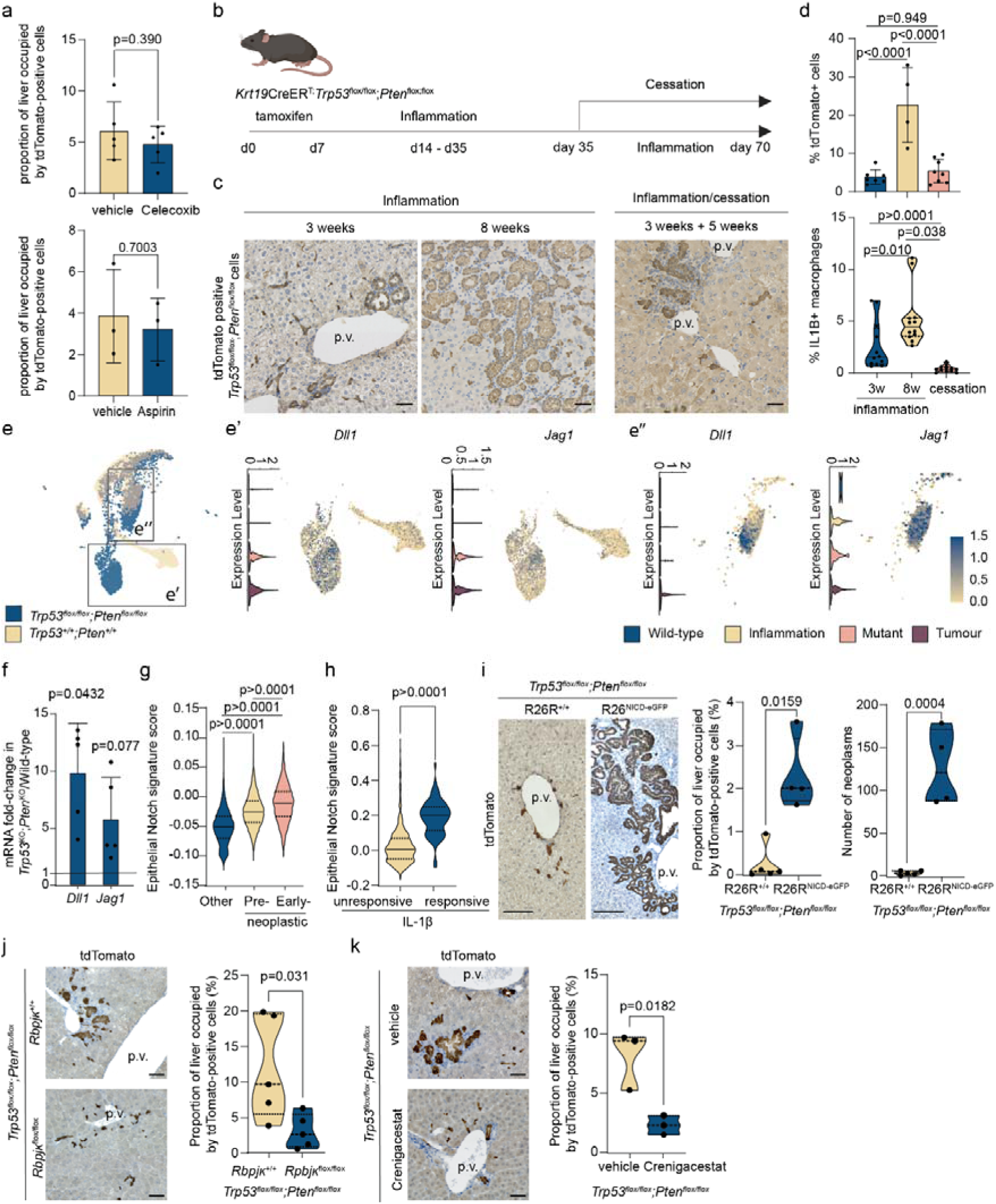
Targeting oncofoetal cell states prevents early-neoplastic transformation. **a.** Quantification of tdTomato-positive tumour number in *Krt19*-CreER^T^;R26R^LSL-tdTomato^;*Trp53*^flox/flox^;*Pten*^flox/flox^ animals treated with vehicle (N=5 individual animals) or Celecoxib (N=5 individual animals), or vehicle (N=3 individual animals) or Aspirin (N=3 individual animals). **b.** Schematic detailing the inflammation treatment and cessation regimen used to treat *Krt19*-CreER^T^;R26R^LSL-tdTomato^;*Trp53*^flox/flox^;*Pten*^flox/flox^. **c.** Immunohistochemistry for tdTomato-positive tumour cells following 3 and 8 weeks of inflammation or 3 weeks, followed by cessation, (Scale bar = 100µm)**. d.** Quantification of tdTomato-positive cells (upper panels, each point represents an individual animal) and IL-1β-positive macrophages (lower panel, each point represents an individual region) in *Krt19*-CreER^T^;R26R^LSL-tdTomato^;*Trp53*^flox/flox^;*Pten*^flox/flox^ animals with 3 or 8 weeks of inflammation or cessation. **e.** UMAP showing *Trp53*;*Pten*-mutant (blue) or wild-type (beige) BECs exposed to inflammation. e’ shows large duct BECs and e’’, early-neoplastic cells and the mRNA expression for *Dll1* and *Jag1*. Histograms show the level of *Dll1* and *Jag1* across experimental groups. **f.** mRNA expression of *Dll1* and *Jag1* in gRNA^*Trp53*^;gRNA^*Pten*^ compared to ICOs transduced with a non-targeting gRNA (used to define the baseline at 1, n=5 experimental replicates). **g.** Expression of transcriptional epithelial Notch signature in murine pre-neoplastic (beige), early neo-plastic (pink) and all other cells (blue). **h.** Transcriptional epithelial Notch signature in human BECs which are IL-1β-unresponsive (beige) and IL-1β-responsive (blue). *i. Krt19*-CreER^T^;R26R^LSL-tdTomato^;*Trp53*^flox/flox^;*Pten*^flox/flox^ mice with R26R^LSL-Nicd:GFP^ or control with no Nicd over expression (R26R^+/+^). Left panels, immunohistochemistry of tdTomato in these lines (scale bar=200µm). Histograms showing proportion of liver occupied by tdTomato mutant cells (left) or number of neoplasms (right) when N^icd^ is overexpressed, (N=4-5 individual animals). *Krt19*-CreER^T^;R26R^LSL-tdTomato^;*Trp53*^flox/flox^;*Pten*^flox/flox^ animals lacking Notch pathway through either **j.** *RBPJκ*^flox/flox^ or through treatment with **k.** Crenigacestat. Immunohistochemistry shows tdTomato-positive *Trp53*;*Pten*-mutant cells (scale bar = 100µ and histograms, proportion of liver occupied by mutant cells (N=3 or 5 individual animals per group).

To test this, we gave *Trp53*;*Pten*-mutant mice inflammation followed by a period of cessation (Fig. 4b) which stalls the progression of neoplastic growth (Fig. 4c and 4d) and is associated with a loss of IL-1β+ macrophages. In fact, regardless of genotype, tissue damage and inflammation through TAA recruits IL-1β-positive macrophages to both peri-central and peri-portal regions of the liver lobule (Extended Data Fig. 5a-c), demonstrating that, in contrast to KRas^G12D^-driven PDAC, the recruitment of IL-1β-positive macrophages is not COX2-dependent when tissue is inflamed. There is, however, a significant increase in peri-portal, BEC-adjacent, IL-1β-positive macrophages in *Trp53*;*Pten*-mutant mice (which is not found in peri-central regions of the lobule), suggesting that mutational status may enhance the ability of COX2-positive, pre-neoplastic cells to recruit pro-tumorigenic macrophages specifically to peri-ductular areas, where tumours arise (Extended Data Fig. 5b and 5c).

## Mutant BECs reactivate canonical developmental programmes to drive neoplastic growth

While targeting COX2 is insufficient to prevent inflammation-induced tumorigenesis, our work shows that the early phases of neoplastic growth require inflammation and are not self-sufficient. Many tumours activate “oncofoetal” programmes to support their growth^33–35^. Notch signalling is an archetypal regulator of bile duct development^36,37^; therefore, we hypothesised that COX2 pre-neoplastic cells acquire an oncofoetal state through reactivation of Notch.

Following *Trp53*;*Pten*-loss and inflammation, both pre- and early-neoplastic cells are specifically enriched for the canonical Notch ligands *Dll1* and *Jag1* (Fig. 4e). The expression of Notch ligands comprises part of the response to *Trp53* and *Pten*-loss as deletion of these genes in otherwise wild- type intrahepatic cholangiocyte organoids (ICOs) using Cas9-CRISPR results in the transcriptional induction of *Dll1* and *Jag1* mRNA (Fig. 4f). Notch is a mitogen in established CCA^38–40^ and therefore we reasoned that the activation of Notch signalling in early neoplasms promotes tumour initiation. Using *Trp53*;*Pten*-mutant mice containing a Notch signalling reporter (RBPJκ-H2B^mVenus^) we show that Notch-responsive mutant BECs are present within early-neoplasms (Extended Data Fig. 6a and 6b). When isolated into Notch active (tdTomato+;mVenus+) or inactive (tdTomato+;mVenus-) cells using FACS (Extended Data Fig. 6c), those with higher levels of Notch activity formed ICOs more rapidly compared to Notch-low counterparts, suggesting that the acquisition of a Notch-high state imparts a progenitor-like phenotype on mutant BECs and drives neoplastic growth (Extended Data Fig. 6d).

To test this, we ectopically expressed the Notch-1 intracellular domain (N^icd^) in *Trp53*;*Pten* transformed BECs (using R26R^LSL-Nicd:GFP^). Ectopic activation of Notch signalling drives increased tumour size and number following inflammation (Extended Data Fig. 6e) and is driven through increased proliferation (Extended Fig. 6f). Bulk RNA sequencing of isolated *Trp53*;*Pten* mutant BECs with either normal levels of Notch activity or overexpressing N^icd^ were used to develop a mutant BEC-specific Notch transcriptional signature (Extended Fig. 6g, Supplementary Table 8). When imposed onto states defined by Stator, this signature identifies a step-wise progression of Notch pathway activation as cells transition to a pre-neoplastic state, which is further enhanced as BECs enter a *Ptgs2*/*Ccl2*/*Mmp7*-positive state (Fig. 4g, Supplementary Table 9). Similarly, IL-1β-responsive BECs from PSC patients are Notch-responsive, showing enrichment for a previously published Notch signature^41^ (Fig. 4h, Supplementary Table 10).

## Inhibition of Notch signalling prevents BilIN formation

Given activation of Notch exaggerates neoplastic transformation, we sought to definitely address whether Notch signalling is sufficient for *Trp53*;*Pten*-mutant BECs to become neoplasms. Overexpression of N^icd^ concurrently with *Trp53*;*Pten*-deletion induced BilIN formation after eight weeks in the absence of inflammation (Fig. 4i), demonstrating that Notch activity supersedes the requirement for inflammation in promoting neoplastic transformation. We rationalised then that mitogenic signalling through Notch actually represents a downstream outcome of inflammation, and unlike COX2-inhibition (where efficacy is dependent on cell-extrinsic inflammation influencing macrophage recruitment), inhibiting the activation of Notch directly would prevent tumour initiation regardless of inflammatory context. To test this, we took two analogous approaches: Firstly, we deleted the essential Notch transcriptional, co-activator, RBPJκ (using *Rbpjκ*^flox/flox^ mice) in *Trp53*;*Pten* mutant cells and second, we treated animals harbouring *Trp53*;*Pten*-loss with the γ-secretase inhibitor Crenigacestat (LY3039478) during the period of inflammation-driven tumour initiation. Genetic deletion of *Rbpjκ* or pharmacological inhibition of Notch signalling almost completely ablated the ability of *Trp53*;*Pten*-mutant BECs to transition into BilIN-like neoplasms (Fig. 4j and 4k), highlighting targeted Notch inhibition as a prophylactic avenue for restricting neoplastic progression in chronic inflammatory disease.

## Discussion

Mutant cells are retained in adult tissues, but do not necessarily contribute to cancer^5,6,11^. Chronic inflammatory disease, however, acts as a potentiator of cancer formation across tissues suggesting that mutant cells might contribute to early neoplastic outgrowth by perceiving inflammatory signals differently to their wild-type neighbours^9^ and by failing to be eliminated through cellular competition^42–44^. Using the mammalian bile duct as an example of a regenerative tissue^45,46^, we demonstrate that cells carrying pathogenic mutations are poised to progress into neoplasia, but only do so when the inflammatory context is permissive, requiring IL-1β-signalling to potentiate an oncofoetal switch that facilitates the formation of a pathological neoplasm.

Beyond demonstrating that inflammation is required for tumour initiation we also show that common, cancer-causing mutations function differently in relatively homogenous cell populations; by using deep single-cell sequencing of BECs, we identify that large duct cells are uniquely sensitive to *Trp53*-loss and PI3K-activation (through deletion of *Pten*), providing experimental support to explain why large duct inflammation in human PSC is associated with increased cancer risk while small duct inflammation in PBC and small duct PSC is not^8,29,30^. Targeted screening has failed to find robust cellular markers of neoplastic transformation in the bile duct, making prediction of likelihood-to-progression in high-risk patients impossible. Using a probabilistic approach to identify cell states from single-cell transcriptomic data^24^, we discovered that a COX2-expressing monocyte recruitment phenotype drives large duct cells into early biliary neoplasms, and show that COX2 demarcates potentially neoplastic cell states in pre-cancerous human disease. Critically, these cells respond to IL-1β analogous to epithelial-macrophage crosstalk previously identified in KRas^G12D^-driven PDAC^9^, suggesting that neoplastic transitions in ductular carcinomas follow a generalisable route through which tumour initiation occurs.

While inflammation promotes the formation of neoplasms, tissue-level injury and inflammation overcomes the requirement for mutant cells to drive local immune recruitment, limiting the effectiveness of targeting immune modulators specifically expressed by dysplastic epithelia, such as COX2 – explaining, perhaps, why clinical trials exploring the use of NSAIDs to reduce cancer risk have had limited sucess^47^. A better understanding of the influence of local inflammatory context, therefore is required when rationalising immunomodulatory or anti-inflammatory approaches to treat patients with chronic disease who are at risk of developing cancer. To this end, we propose in lieu of targeting inflammation, a more reasoned approach is to target the essential oncofoetal signals which neoplasia rely on to form and which are able to modulate their local inflammatory milieu^33,34^. Indeed, we show that pre- and early-neoplastic cell states reactivate the archetypal biliary developmental signal, Notch^48,49^ and by preventing this oncofoetal switch, we prevent tumour initiation. Collectively, our data maps the earliest phases of tumour initiation within the bile duct and shows that this process can be reversed – highlighting a route to prophylactic treatment in high-risk patients.

## Methods

### Animal experiments

All *in vivo* experiments were carried out in accordance with the guidance issued by the Medical Research Council in “Responsibility in the Use of Animals in Medical Research” (July 1993) and licensed by the Home Office under the Animals (Scientific Procedures) Act 1986. Experiments were performed under project license number PFD31D3D4 in facilities at the University of Edinburgh (PEL 60/6025). Animals were housed with food ad libitum in 12h light-dark cycles.

For modelling inflammation-dependent tumour initiation in mice we used previously described^21^ compound mutant line where Keratin-19-CreER^T^ (Jax: 026925);*Trp53*^flox/flox^ (Jax: 008462);*Pten*^flox/flox^ (Jax: 006440);R26R^LSL-tdTomato^(Jax: 007908) model or as a control Keratin-19-CreER^T^; R26R;*Trp53*^+/+^ (Jax: 008462);*Pten*^+/+^. For the induction of genetic recombination and liver inflammation in mice, 4mg of tamoxifen resuspended in 90% Corn oil, 10% ethanol was given by oral gavage three times on alternate days. After one-week mice were provided with thioacetamide (TAA)(400mg/L) *ad libitum* in sweetened drinking water. In addition to this baseline model, Notch-activity reporting and over-expression lines were generated by crossing in Tg(Cp-HIST1H2BB/Venus)47Hadj/J (CBF-H2B:mVenus, #020942) or Gt(ROSA)26Sor^tm1(Notch1)Dam^/J (Jax: 008159), respectively.

For therapeutic intervention experiments mice were treated with either; Celecoxib given daily Mon-Fri as a 400 µg oral gavage (resuspended in 40% H_2_O, 50% PEG400, 10% DMSO), aspirin provided *ad libitum* at 1200 mg/L in TAA drinking water, or crenigacestat (LY3039478) at 8mg/Kg, three-times weekly by oral gavage.

### Histopathology and immunofluorescence

Mouse livers were perfused with phosphate buffered saline and fixed overnight in 10% neutral buffered formalin. Following standard processing tissues were sectioned at 5µm. Sections were immunostained using the antibodies detailed in Supplementary Table 11. Immunofluorescent slides were mounted in Vectashield mountant containing DAPI (VectorLabs). For colorimetric staining biotinylated secondary antibodies were used, detailed in Supplementary Table 11 and were counterstained with haematoxylin. Slides were imaged using a Nanozoomer slide scanner (Hamamatsu), and immunofluorescent staining was imaged using the Zeiss Axioscan.Z1 slide scanner or a Nikon A1R confocal microscope. Shading correction was performed on fluorescent whole slide images to account for tiling effects using Zeiss Zen 3.5 software. Tissue clearing for full duct imaging was performed using a modified FUnGI protocol as previously described^21,50^. Briefly, cores of liver tissue were sectioned into 200µm thick sections using a Krumdieck Tissue Slicer, fixed for 45 min in 10% neutral buffered formalin before being washed overnight at 4°C (PBS, 0.1% Tween20, 50 µg/ml ascorbic acid, 0.05ng/ml L-Glutathionine reduced) and cleared overnight at 4°C in FUnGI clearing solution (50% glycerol (vol/vol), 2.5 M fructose, 2.5 M urea, 10.6 mM Tris Base, 1 mM EDTA) and mounted and imaged on a Nikon A1R confocal microscope.

### Human tissue

Anonymized patient tissue was acquired with the approval of the Lothian NRS BioResource (SR1872 and SR2185). Patient information is available in Supplementary Table 12.

### Quantification of immunohistochemistry and immunofluorescence

All quantification was performed using the digital pathology software QuPath v 0.5.1^51^. For quantification of tdTomato+ cells, automated cell detection was performed based on haemotoxylin counterstaining of the entire left lateral lobe. Cell positivity was defined through DAB intensity and the proportional positivity determined. For lobe burden; pixel thresholding based on DAB intensity was used to determine proportional area of the lobe stained for tdTomato. For immunofluorescent quantification of IL-1β+ macrophages F4/80 staining was used to detect all macrophages within a 300 µm radius of portal triads or central veins (n=10 regions across n=3 mice). F4/80-positive cells were thresholded for IL-1β based on fluorescent intensity normalised to background intensity to account for autofluorescence. For COX2 scoring in patient tissue, n=5 randomly sampled 20 mm^2^ regions were applied per slide. Epithelial regions were detected using a multiparameter trained classifier based on panCK positivity, and COX2 positive cells were detected based on a positivity threshold normalised to background autofluorescence. For the detection of IL-1β-positive macrophages, total CD68-positive cells were detected in 1 mm^2^ regions (n=5). Within the CD68-positive compartment, IL-1β-positive cells were detected based on fluorescent intensity adjusted for background autofluorescence.

### Digestion and isolated bile ducts

Isolation of murine biliary epithelial cells for downstream applications was performed as follows; dissected liver was finely chopped into 1-3 mm pieces and parenchyma was digested by incubating at 37°C and 150 rpm for 2hrs in Advanced DMEM-F12 (Gibco) supplemented with 1% foetal bovine serum, 1x Antibiotic-antimycotic (Gibco), 1x Glutamax (Gibco), 0.125 mg/ml collagenase-IV (Gibco 17104-019) and 0.125 mg/ml dispase (Gibco 1 17105-041). After parenchymal digestion, non-parenchymal cells were pelleted and washed before resuspending and agitating in 7x TryplE solution to produce a single cell solution. These were filtered through a 40 µm strainer and red blood cells lysed by incubating with ACK lysis buffer (Gibco), and cell pellets resuspended in DMEM-F12 with 1% FCS, 1x Glutamax and 1x Antibiotic-antimycotic ready for flow cytometry.

### Murine intrahepatic cholangiocyte organoids (ICOs)

Murine ICOs were generated from C57/Bl6 mice. Isolated biliary epithelial cells were grown in Matrigel (Corning) in base media comprising Advanced DMEM/F-12 (Gibco) supplemented with 1x Glutamax (Gibco), 1x Antibiotic-antimycotic (Gibco), 1x Fungizone (Gibco), HEPES (Sigma-Aldrich), mEGF (R&D Systems; 50 ng/ml), mFGF-10 (ThermoFisher;100 ng/ml), mHGF (ThermoFisher;5 ng/ml), Gastrin (Sigma-Aldrich;10 nM), Nicotinamide (Sigma-Aldrich; 10 mM), N-acetylcystine (Sigma-Aldrich; 1.25 mM), 1x B27 (Gibco), Y-27632 (Tocris; 10 nM), forskolin (Tocris; 10 nM), A83-01 (Sigma-Aldrich; 5 nM), Chir99021 (ApexBio; 3.3 nM), and passaged twice a week as previously published^52^.

### Generation of *Trp53;Pten* knockout organoids

*Trp53*;*Pten* knock-out organoid lines were generated using lentiviral transduction of CRISPR-Cas9 and pooled guide RNAs targeting the *Trp53* or *Pten* gene. gRNAs against *Trp53* and *Pten* were cloned into Addgene #98290 and Addgene #98293 respectively. Lentiviral particles were generated by transient transfection of these plasmids and packages vectors VSVG (Addgene #8454) and psPax2 (Addgene #12260) into HEK293T cells and isolated from the media after 3 days. Wild-type C57/Bl6 biliary organoids were transduced with viral particles cultured in organoid media supplemented with 2ug/ml puromycin and 10ug/ml blasticidin.

### CBF-H2B:mVenus organoid growth

Bile ducts were isolated from Keratin-19-CreER^T^;*Trp53*^flox/flox^;*Pten*^flox/flox^;R26R^LSL-tdTomato^;CBF-H2B:mVenus reporter mice. Live (DAPI-negative) BECs were sorted based on tdTomato-positivity and gated for high versus low mVenus fluorescence. Organoid area over time was measured using the Incucyte Live-Cell Analysis Instrument (SARTORIUS).

### Bulk RNA sequencing

Total-RNA samples were fragmented to a size appropriate for sequencing on an Illumina platform and first-strand cDNA was generated using the SMARTer® Stranded Total RNA-Seq Kit v2 – Pico Input Mammalian kit (Clontech Laboratories, Inc. #634411). Illumina-compatible adapters and indexes were then added via 5 cycles of PCR. Depletion of ribosomal cDNA (cDNA fragments originating from highly abundant rRNA molecules) was performed using ZapR v2 and R-probes v2 specific to mammalian ribosomal RNA and human mitochondrial rRNA. Sequencing was performed on the NextSeq 2000 platform (Illumina Inc, #20038897) using NextSeq 2000 P2 Reagents (200 Cycles) (#20046812). Libraries were combined in a single equimolar pool of nine based on Qubit and Bioanalyser assay results and run on a single P2 flow cell. PhiX Control v3 (Illumina, #FC-110-3001) was spiked in at 1% library concentration to facilitate troubleshooting in the event of any run issues.

### RNA sequencing data processing and analysis

The primary RNA-Seq processing, quality control to transcript-level quantitation, was carried out using nf-core/rnaseq v1.4.3dev (https://github.com/ameynert/rnaseq)^53^. Reads were mapped to the mouse FVB_NJ_v1 decoy-aware transcriptome using the salmon aligner (1.1.0). RNA-Seq analysis was performed in R (4.0.2), Reads were summarized to gene-level and differential expression analysis was performed using the bioconductor packages tximport (1.16.1) and DESeq2 (1.28.1). A pre-filtering was applied to keep only genes that have at least 10 reads in a group and 15 reads in total. The Wald test was used for hypothesis testing for pairwise group analysis. A shrunken log2 fold changes (LFC) was also computed for each comparison using the adaptive shrinkage estimator from the ’ashr’ package. Bulk RNA sequencing is deposited at GEO, accession numbers: XXXXXXXXX

### Data generation for scRNA-seq

Single cell RNA sequencing (scRNA-seq) of murine biliary epithelial cells was performed using the 10x Genomics Chromium Next GEM 3’ kit. Briefly, tdTomato-tagged neoplasms baring animals and control mice had their livers dissected and ducts isolated (as above). Cells were sorted DAPI negative, tdTomato positive cells using the BD FACSARIA II cell sorter. Single cell GEMs were generated through the 10x microfluidics platform and resulting Chromium 3’ gene expression libraries sequenced using the NextSeq 2000 platform (Illumina) on P3 flow cells. Single cell sequencing data for wild-type animals was integrated from our previous study (Dryad: doi:10.5061/dryad.mkkwh7152) during data processing.

### scRNA-seq data processing

Raw sequencing reads were processed using the 10x Genomics CellRanger pipeline (v.5.0.0) using mouse reference genome (mm10). Resulting gene expression matrices were analysed using the R package Seurat (v.5). Seurat v5.1.0 was used with the exception of marker detection in supplementary figure 1 which used v 4.1.0. Cells were filtered to ensure they had: UMI >650 & <75000, with >300 & <7000 unique genes, <5% mitochondrial gene fraction >0 dtTomato expression and >0.8 log10GenesPerUMI. Empty droplets were removed and doublet detection and removal was performed using Scrublet. Filtered matrices were normalised and data integration performed, Louvain clustering and differential gene expression analysis between clusters was performed in Seurat. Original data can be accessed from DOI: 10.5061/dryad.9kd51c5w1.

### Reanalysis of single cell data from Andrews et al. 2024

Human 5’ scRNA-seq raw count matrices were downloaded from the GEO repository (accession numbers: GSE243977 and GSE247128) and processed using Seurat (v.5) in RStudio using R (v.4.3.2). Cells were filtered to only include those with unique gene counts > 1000 & <4000 and <25% mitochondrial gene fraction. After normalisation and identification of the top 2000 variable features datasets were integrated using the FindIntegrationAnchors and integrateDate functions in Seurat^54^ before following the standard Seurat pipeline for PCA, heirarchical clustering and UMAP projection.

### RNA extraction

RNA was extracted using TRIzol RNA Isolation Reagent (Invitrogen), precipitated with chloroform, and cleaned up using the RNeasy Mini Kit (Qiagen). Quantiect® reverse transcriptase kit (QIAGEN), including gDNA Wipeout buffer for elimination of genomic DNA, was used as per manufacturer’s instructions for cDNA synthesis. Quantitative PCR (qPCR) was performed with SYBR™ Green PCR Master Mix (Applied Biosystems™), and real-time PCR run using the Roche LightCycler® 480 II. For downstream sequencing applications RNA quality (RIN score) was quantified using the Agilent 2100 Bioanalyzer with an RNA 6000 chip. QRTPCR was performed using primers in Supplementary Table 13.

### Usage of additional R packages

Differential neighbourhood abundance testing was performed using the MiloR package^23^ (v 2.0.0). From Seurat objects, k nearest neighbour graphs were created using the buildGraph function (k=10)(d=30), representative cell neighbourhoods were then generated using makeNhoods (prop=0.1). Differential abundance of groups within representative neighbourhoods was then tested using the testNhoods function to provide log fold change and spatially corrected p-value for each neighbourhood. To generate a list of top 50 differential expressed genes we used the findNhoodGroupMarkers function specifically within Seurat cluster 4.

For ORA and GSEA analysis of GO terms was performed using clusterProfiler (v. 4.12.6). ORA within gene sets was performed for gene ontology terms associated with biological process (BP) using the enrichGO function. GSEA was performed using the gseDO and plots were generated using gseaplot2. Tree plots were used to heirarchically cluster enriched terms based on pairwise similarities determined using the Jaccard’s similarity index and were generated using the treeplot function. Finally, Manhattan plots for wider GO assessment were created using the gostplot function in gprofiler2 (v. 0.2.3).

### Stator defined cell states from higher-order gene expression dependencies

We applied Stator to identify cell states. Briefly, Stator initially takes in binarised scRNA-seq data and estimates higher-order interactions (HOIs) among top highly variable genes. Gene tuples driving these HOIs were then extracted by testing from the null hypothesis of independence with log2-transformed deviating enrichment factor of 1.5, and termed D-tuples. Hierarchical clustering of D-tuples was then performed to group these D-tuples, with dice distance of 0.76. To explore whether there are any Stator states are more enriched in a specific condition, we performed hypergeometric test for the enrichment of a Stator states in different conditions. We then used the Benjamini-Hochberg (BH) procedure for multiple testing correction.

### Statistical analysis

Statistical analysis was performed using GraphPad Prism 9. Exact P values are noted in the figure. All data was tested for normality using a D’Agostino-Pearson test for normality. In cases of two group comparisons t-tests (or non-parametric alternatives) were used. For multigroup testing (>2) ANOVA with Tukey HSD *post-hoc* was used.

## Acknowledgements

We would like to thank the Institute of Genetics and Cancer Advanced Imaging Resource for their support with imaging and the IGC Cytometry and Single Cell Core facility for support with flow cytometry analysis and cell sorting. We would also like to thank staff at the University of Edinburgh Bioresearch & Veterinary Services for husbandry support.

## Author Contributions

EJJ developed concepts, performed experiments, generated and analysed data, drafted the manuscript and provided financial support for the study. AML developed concepts, performed experiments, generated and analysed data. YY analysed and generated data for the project. AR, SW, AG generated and analysed data, KD provided technical assistance with *in vivo* work. RG provided conceptual support. SR provided tissue and conceptual support. TJK and OJ provided resources/samples and analytical input into the project. AK provided academic direction to project, developed and supported methodological advances, analysed data and edited the manuscript. LB lead and conceptualised the project, provided guidance and intellectual input into the concepts, analysed data, edited and finalised the manuscript and provided funding for the project.

## Data and materials availability

All data is available in the manuscript or the supplementary materials. Previously published single cell RNAseq data from this study is available from: GSE243981. Data generated in this study is available at Dryad: DOI: 10.5061/dryad.9kd51c5w1. All materials generated as part of this study will be made available upon request to the corresponding authors.44

## List of Extended data figures and Tables

Extended Data 1: Initiating cancer through inflammation and tumour suppressor-loss

Extended Data 2: Transformed cells have a unique transcriptional profile independent of inflammation.

Extended Data 3: Neoplasms have a specific transcriptional cell state:

Extended Data 4: IL-1β-positive macrophages are physically associated with neoplastic cells.

Extended Data 5: Single cell RNA sequencing identifies disease specific-cellular relationships in high-risk patients.

Extended Data 6: Notch activation drives transformed cells into neoplasms.

## List of Supplementary Tables

Supplementary Table 1: Seurat clusters marker table

Supplementary Table 2: Common gene lists for inflammation and mutation

Supplementary Table 3: Large duct DEG analysis

Supplementary Table 4: Regenerative cluster neighbourhood DEG analysis

Supplementary Table 5: Stator transition DEG analysis

Supplementary Table 6: IL1β epithelial response signature genes

Supplementary Table 7: Receptor ligand analysis

Supplementary Table 8: DEG normalised read counts following Notch overexpression

Supplementary Table 9: Murine Notch biliary gene signature

Supplementary Table 10: Human Notch signature and scoring (Vilimas et al.)

Supplementary Table 11: Antibodies used in this study

Supplementary Table 12: Human Tissue used within this study

Supplementary Table 13: Oligos used in this study

