## Supplementary Figures for "Epithelial state-transitions permit inflammation-induced tumorigenesis"

**List of Extended Data Figures:**

Extended Data 1: Initiating cancer through inflammation and tumour suppressor-loss

Extended Data 2: Transformed cells have a unique transcriptional profile independent of inflammation.

Extended Data 3: Neoplasms have a specific transcriptional cell state:

Extended data 4: Single cell RNA sequencing identifies disease specific-cellular relationships in high-risk patients.

Extended data 5: IL-1β-positive macrophages are physically associated with neoplastic cells.

Extended Data 6: Notch activation drives transformed cells into neoplasms.

**List of Supplementary Tables:**

Supplementary Table 1:	Seurat clusters marker table

Supplementary Table 2:	Common gene lists for inflammation and mutation

Supplementary Table 3:	Large duct DEG analysis

Supplementary Table 4:	Regenerative cluster neighbourhood DEG analysis

Supplementary Table 5:	Stator transition DEG analysis

Supplementary Table 6:	IL1β epithelial response signature genes

Supplementary Table 7:	Receptor ligand analysis

Supplementary Table 8:	DEG normalised read counts following Notch overexpression

Supplementary Table 9:	Murine Notch biliary gene signature

Supplementary Table 10: Human Notch signature and scoring (Vilimas et al.)

Supplementary Table 11: Antibodies used in this study

Supplementary Table 12: Human Tissue used within this study

Supplementary Table 13: Oligos used in this study

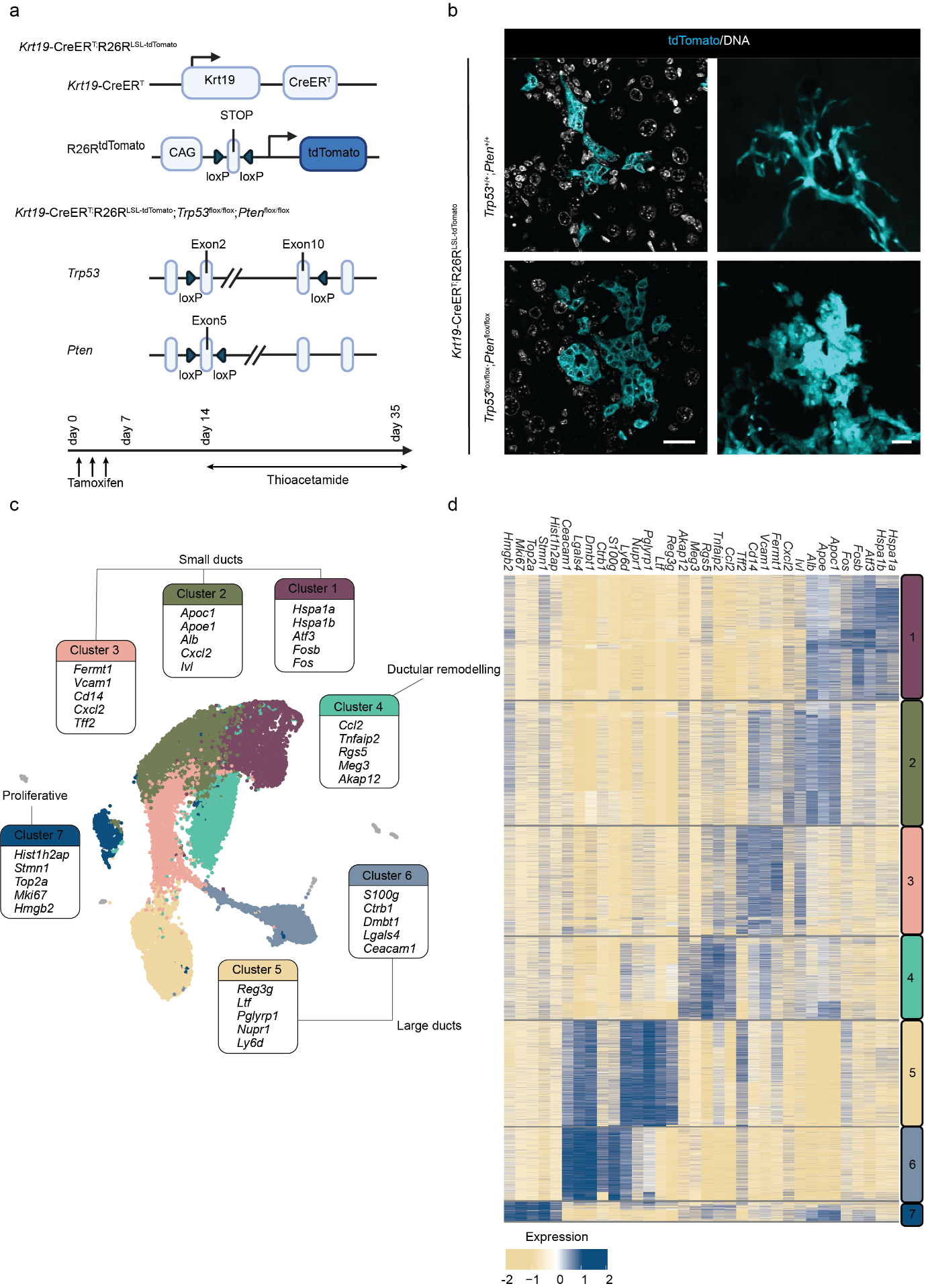

**Extended Data 1: Initiating cancer through inflammation and tumour suppressor-loss:** **a.** Schematic detailing the approach by which *Trp53* and *Pten* are deleted selectively within biliary epithelial cells (BECs) while irreversibly labelled with the bright fluorophore, tdTomato. **b.** Immunofluorescent staining of tdTomato (cyan) and DNA (white) in either *Krt19*-CreERT;R26RtdTomato;*Trp53*^+/+^;*Pten*^+/+^ or *Krt19*-CreERT;R26RtdTomato;*Trp53*^flox/flox^;*Pten*^flox/flox^. Scale bar = 100µm**.** **c.** Seurat clustering of fluorescently sorted tdTomato-positive BECs with and without *Trp53*;*Pten*-loss and with or without inflammation. **d.** Heatmap showing differentially expressed genes between the 7 principle Seurat groups.

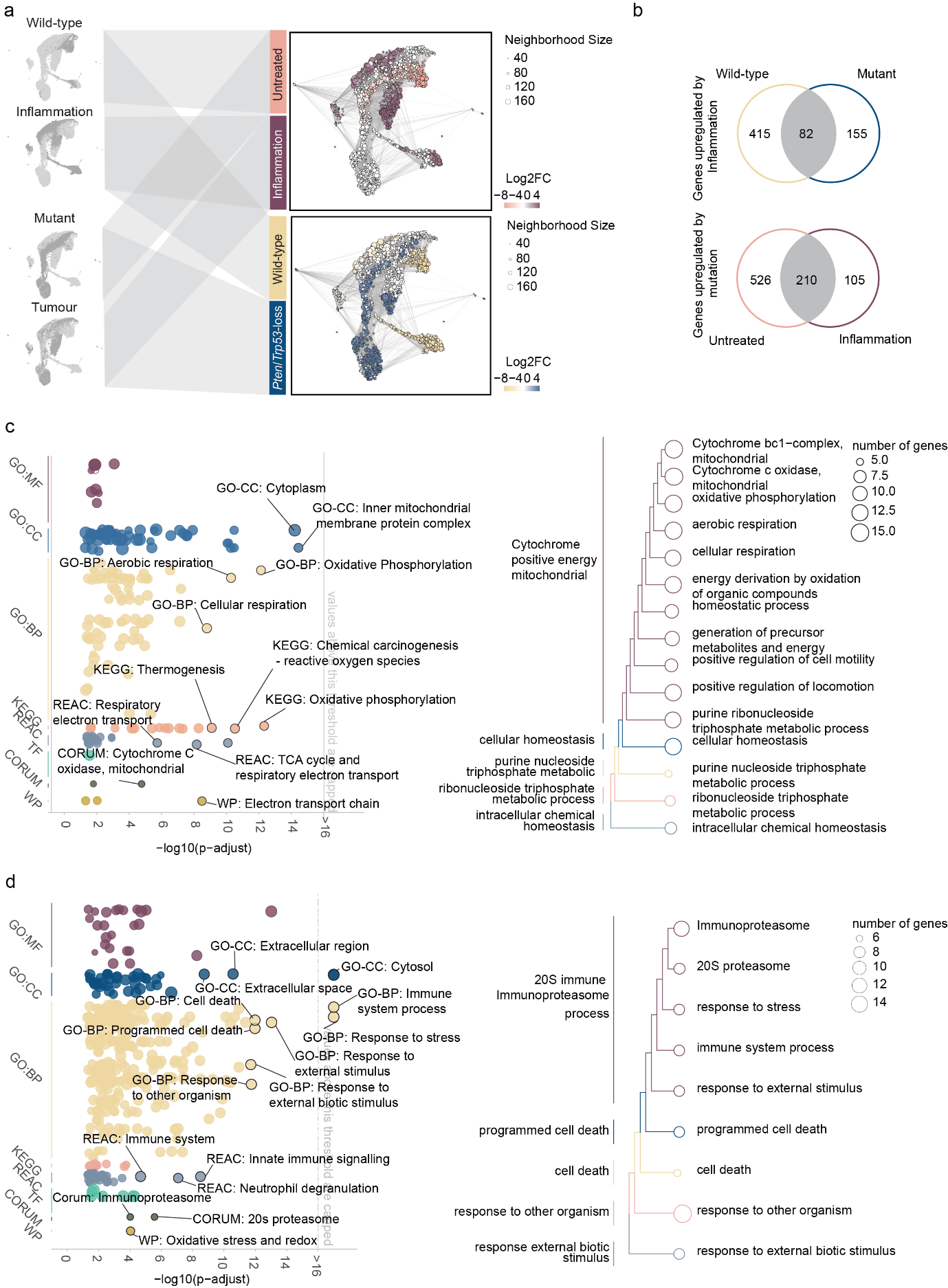

**Extended Data 2: Transformed cells have a unique transcriptional profile independent of inflammation. a.** Milo generated cellular neighbourhoods which arise in single cell data from *Trp53*;*Pten*-mutant BECs with and without inflammation. Neighbourhoods showing differential abundance were identified with SpatialFDR < 5%. **b.** Venn diagram demonstrating the number of genes up- or down- regulated following mutation and/or inflammation in single cell data. **c.** Functional enrichment analysis of genes upregulated in response to inflammation represented as a Manhattan plot of enriched terms (left panel) and Treeplot showing hierarchical clustering of enriched terms (right panel) **d**. Manhattan (left panel) and Treeplot (right panel) of functional enrichment analysis from genes upregulated by *Trp53*:*Pten* deletion.

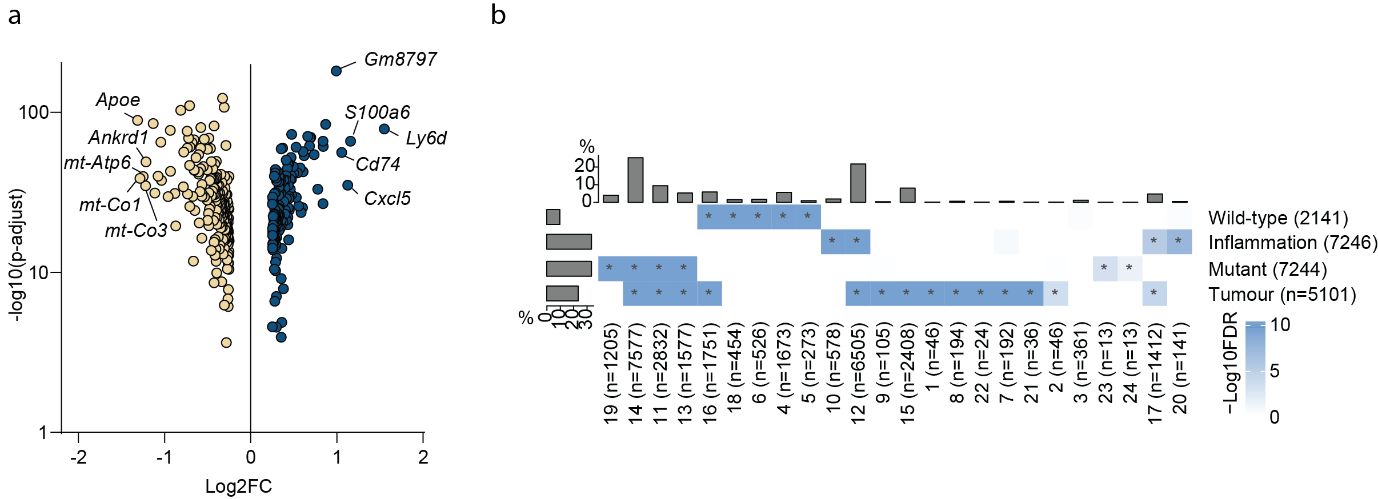

**Extended Data 3: Neoplasms have a specific transcriptional cell state: a.** Volcano plot demonstrating the up- and down-regulated genes from neoplastic cell neighbourhoods in cluster 4. **b.** Stator cell state outcomes from wild-type, inflammation, mutant and tumour groups. The x-axis represents Stator states, and *n* denotes number of cells in each Stator state. Blue boxes with an asterisk represent Stator states that are significantly enriched in each of the four conditions (FDR < 0.05).

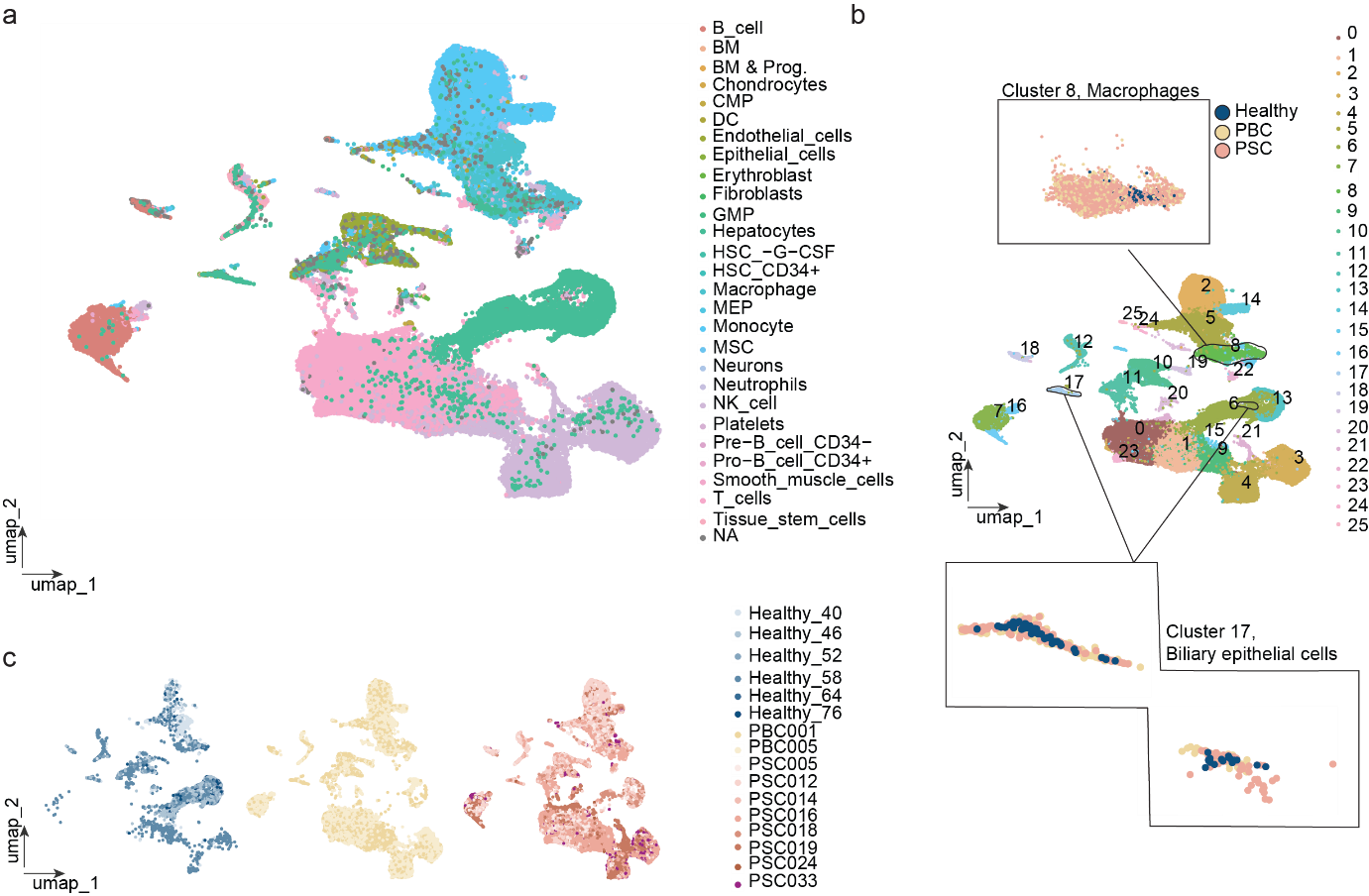

**Extended data 4: Single cell RNA sequencing identifies disease specific-cellular relationships in high-risk patients**. **a.** SingleR analysis of healthy, PBC and PSC patients identifies multiple cell populations within tissue samples. **b.** Seurat clustering of human patient samples identifying 25 independent clusters. Insets show the composition of cells within cluster 8 and cluster 17, with healthy denoted in blue, PBC; beige and PSC; pink. **c.** UMAP projection of healthy, PSC and PBC single cell data separated by group and coloured based on patient sample.

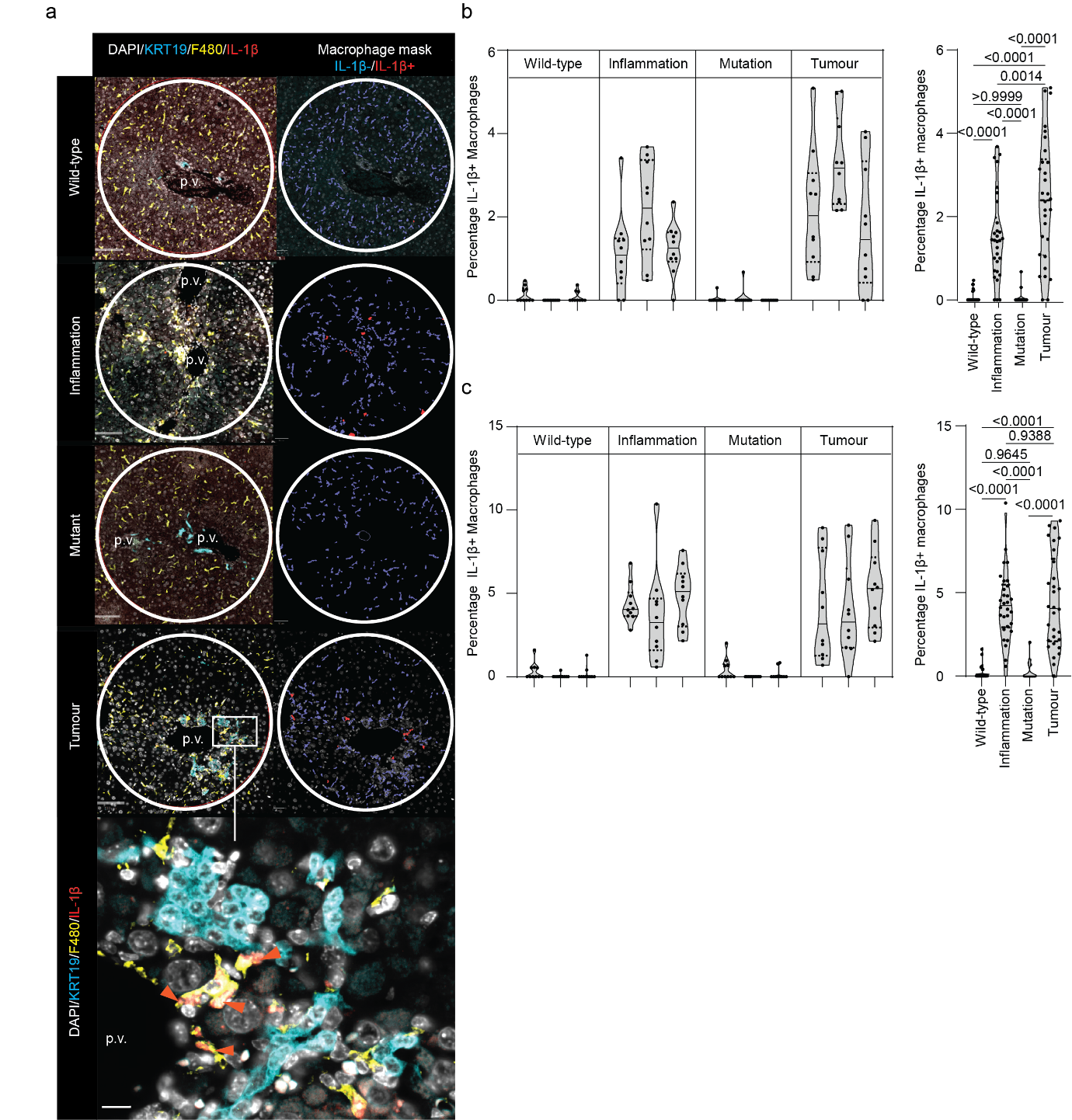

**Extended data 5: IL-1β-positive macrophages are physically associated with neoplastic cells.** **a.** Immunofluorescent staining of KRT19-positive BECs (cyan), F4/80-positive macrophages (yellow) and IL-1β (red), DNA (white) in *Trp53*;*Pten*-mutant or wild-type animals with or without inflammation. **b.** Quantification of IL-1β-positive macrophages in peri-central and **c.** periportal areas**)** in *Trp53*;*Pten*-mutant or wild-type animals with or without inflammation. Scale bar = 200µm, lower panel 25µm.

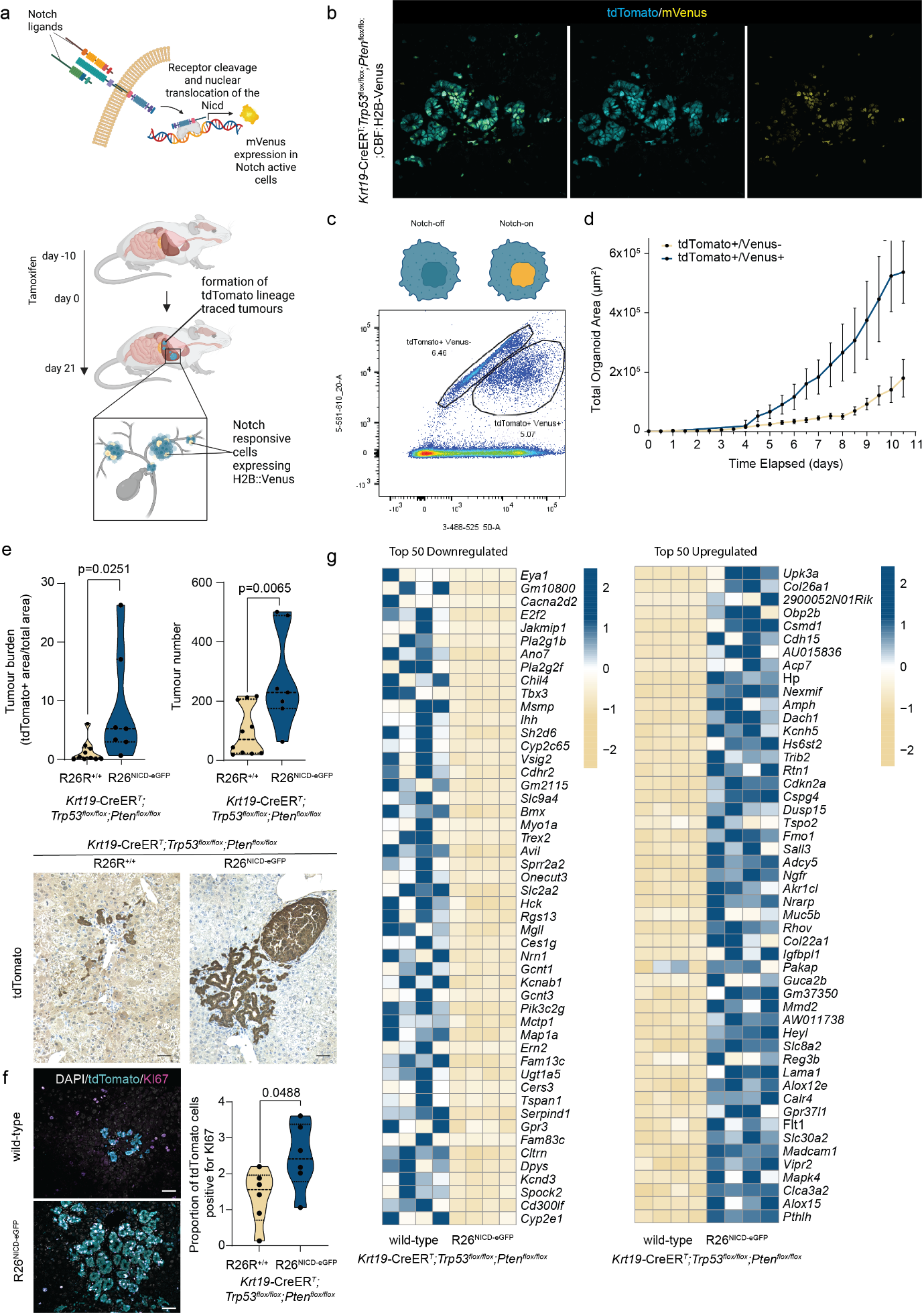

**Extended Data 6: Notch activation drives transformed cells into neoplasms.** **a.** Schematic demonstrating how *Krt19*-CreER^T^;*Trp53*^flox/flox^;*Pten*^flox/flox^;R26R^LSL-tdTomato^;*CBF:H2B*^mVenus^ can be used to identify Notch-responsive cells in early biliary neoplasms. **b.** Immunofluorescence showing a neoplasm from *Krt19*-CreER^T^;*Trp53*^flox/flox^;*Pten*^flox/flox^;R26R^LSL-tdTomato^;*RBPJκ*^mVenus^ mice stained for with tdTomato (cyan) and mVenus (yellow) scale bar = 100µm. **c.** Representative FACS plot showing TdTomato+/mVenus- and TdTomato+/mVenus+ gating strategies. **d.** Total organoid area when FACS isolated cells are plated as organoids from Tomato+/mVenus- (yellow line) and TdTomato+/mVenus+ (blue line) n = 3 biological replicates. **e.** Area of the liver occupied by tdTomato-positive tumour cells (left) and tumour number (right) in *Krt19*-CreER^T^;*Trp53*^flox/flox^;*Pten*^flox/flox^;R26R^LSL-tdTomato^;R26R^LSL-Nicd^ (N=8 individual animals) or R26R^+/+^ (N=9 individual animals) given three weeks of inflammation**.** Lower panels, immunohistochemistry for tdTomato (brown) in these lines. Scale bar = 200µm. **f.** Immunofluorescent staining for tdTomato (cyan), Ki67 (magenta) and DNA (white) of neoplasms either expressing R26R^LSL-Nicd^ or not. Scale bar = 150µm. Violin plot shows the number of tdTomato-positive *Trp53*;*Pten-*mutant cells that are positive for Ki67 **g.** Heatmaps showing the top 50 Up- and Down-regulated genes from FACS isolated neoplastic cells overexpressing Nicd vs those which are wild-type for the Notch intracellular domain.
